# Chemical variations in *Quercus* pollen as a tool for taxonomic identification: implications for long-term ecological and biogeographical research

**DOI:** 10.1101/761148

**Authors:** Florian Muthreich, Boris Zimmermann, H. John B. Birks, Carlos M. Vila-Viçosa, Alistair W.R Seddon

## Abstract

vi.

**Aim:** Fossil pollen is an important tool for understanding biogeographic patterns in the past, but the taxonomic resolution of the fossil-pollen record may be limited to genus or even family level. Chemical analysis of pollen grains has the potential to increase the taxonomic resolution of pollen, but present-day chemical variability is poorly understood. This study aims to investigate whether a phylogenetic signal is present in the chemical variations of *Quercus* L. pollen and to assess the prospects of chemical techniques for identification in biogeographic research.

**Location:** Portugal

**Taxon:** Six taxa (five species, one subspecies) of *Quercus* L., *Q. faginea, Q. robur, Q. robur* ssp. *estremadurensis, Q. coccifera, Q. rotundifolia* and *Q. suber* belonging to three sections: *Cerris, Ilex*, and *Quercus* (Denk, Grimm, Manos, Deng, & Hipp, 2017)

**Methods:** We collected pollen samples from 297 individual *Quercus* trees across a 4° (∼450 km) latitudinal gradient and determined chemical differences using Fourier-transform infrared spectroscopy (FTIR). We used canonical powered partial least-squares regression (CPPLS) and discriminant analysis to describe within- and between-species chemical variability.

**Results:** We find clear differences in the FTIR spectra from *Quercus* pollen at the section level (*Cerris*: ∼98%; *Ilex*: ∼100%; *Quercus*: ∼97%). Successful discrimination is based on spectral signals related to lipids and sporopollenins. However, discrimination of species within individual *Quercus* sections is more difficult: overall, species recall is ∼76% and species misidentifications within sections lie between 18% and 31% of the test-set.

**Main Conclusions:** Our results demonstrate that subgenus level differentiation of *Quercus* pollen is possible using FTIR methods, with successful classification at the section level. This indicates that operator-independent FTIR approaches can surpass traditional morphological techniques using the light microscope. Our results have implications both for providing new insights into past colonisation pathways of *Quercus*, and likewise for forecasting future responses to climate change. However, before FTIR techniques can be applied more broadly across palaeoecology and biogeography, our results also highlight a number of research challenges that still need to be addressed, including developing sporopollenin-specific taxonomic discriminators and determining a more complete understanding of the effects of environmental variation on pollen-chemical signatures in *Quercus*.

## 1. Introduction

Sub-fossil pollen remains preserved in lake sediments or peat bogs have been important tools to reconstruct past floristic, vegetational, and environmental changes for over one hundred years. The biogeographic applications of such reconstructions are varied and wide-ranging. Palaeoecological studies based on fossil pollen have made vital contributions to understanding the broad-scale range dynamics through time, the rates and directions of spread of different plant species, and the location of glacial-stage refugia (see Birks, 2019 for a review). Fossil pollen data can also be used to track relative-niche shifts in association with the emergence of no-analogue climates (Veloz et al., 2012) and forecast future range shifts as a result of climate change (e.g. Nogués-Bravo et al., 2016, 2018).

The basis of all such studies is reliable identifications of fossil pollen to the lowest taxonomic level possible. With detailed identifications, reconstructions and answers to particular biogeographical and ecological questions can similarly be detailed. Indeed, many advances in historical plant geography (e.g. Godwin, 1975; Lang, 1994; Magri et al., 2006; Birks, 2008; Birks, 2014) have been made because of advances in the identification of plant fossils. However, although Quaternary botany (sensu Birks, 2019) has been dominated for over one hundred years by pollen analysis, identifications can only be made to the genus or family level for many taxa. This is limiting the biogeographical information gained from fossil pollen to relatively coarse taxonomic levels.

This issue is of particular relevance for understanding the past, present and future distributions of the genus *Quercus* (oak). *Quercus* contains 22 native species in two subgenera and three sections in Europe (Tutin et al., 1993; Denk et al., 2017), several of which have striking and often distinct geographical distributions today (e.g. Iberia, Balkans, eastern Mediterranean, widespread Mediterranean, Apennine Peninsula, widespread to about 60°N; Jalas & Suominen, 1976). However, what is known about the history of *Quercus* is almost entirely based on pollen and is thus only at the genus level. Although three pollen-morphological types can, with care, be distinguished by conventional light microscopy (LM) (Beug, 2004) and scanning election microscopy (SEM) (Denk & Grimm, 2009), fossil pollen of *Quercus* is usually determined as either *Quercus* Deciduous or *Quercus* Evergreen types.

This situation has several implications for biogeographic research. Maps of the changing distribution and abundance of oak pollen in the late-glacial and Holocene of Europe (Huntley & Birks, 1983; Simon Brewer et al., 2017) can only confidently be made using the two broad pollen morphotypes (i.e. *Quercus* Deciduous or *Quercus* Evergreen). Palaeo-biomisation methods used to forecast the future responses of Mediterranean ecosystems to climate change have used the same distinction between these two morphotypes (Guiot & Cramer, 2016), whilst a recent attempt to model future responses of *Quercus* responses in Europe using fossil pollen were based on *Quercus* pollen resolved to the genus level (Nogués-Bravo et al., 2016). Since the sensitivity and response of *Quercus* to recent environmental changes is species-specific in Mediterranean ecosystems (Acácio, Dias, Catry, Rocha, & Moreira, 2017), and since *Quercus* macrofossils are very rarely found, any improved understanding of its historical and future biogeography clearly depends on consistent pollen identifications at levels lower than is presently available.

One potential approach lies in the chemical analysis of pollen. Fourier Transform Infrared Spectroscopy (FTIR) is a non-destructive method which is used to infer the chemical composition of a sample based on the fact that different molecular-functional groups have different wavelength-specific absorbances of infrared radiation (Pappas, Tarantilis, Harizanis, & Polissiou, 2003; Ivleva, Niessner, & Panne, 2005; Gottardini, Rossi, Cristofolini, & Benedetti, 2007; Schulte, Lingott, Panne, & Kneipp, 2008; Zimmermann, 2010; Parodi, Dickerson, & Cloud, 2013; Zimmermann & Kohler, 2014; Bağcioğlu, Zimmermann, & Kohler, 2015). Evidence suggests that the analysis of pollen using FTIR may be a useful tool for differentiating between pollen types extracted from sediment sequences (Julier et al., 2016; Woutersen et al., 2018; Jardine, Gosling, Lomax, Julier, & Fraser, 2019), because pollen-grain chemistry may show biogeographical patterns related to phylogeny and environmental conditions (Depciuch, Kasprzyk, Roga, & Parlinska-Wojtan, 2016; Bağcıoğlu, Kohler, Seifert, Kneipp, & Zimmermann, 2017; Zimmermann et al., 2017). However, although taxonomic differentiation of pollen based on the chemical variations inferred by FTIR shows considerable promise, widespread application of FTIR in biogeography and palaeoecology remains limited. This is because lipids and proteins can be major discriminants of variation in FTIR spectra (Bağcioğlu et al., 2017; Zimmermann & Kohler, 2014), but these may not be preserved in sediment sequences alongside the sporopollenin-based exines (Zimmermann, Tkalčec, Mešić, & Kohler, 2015). Therefore the relative importance of the different chemical structures (e.g. lipids, proteins, sporopollenins) that are responsible for discrimination between many modern pollen types still needs to be established. Moreover, influence of various abiotic stressors on pollen chemistry can hinder taxonomic differentiation of pollen samples by FTIR (Lahlali et al., 2014; Depciuch et al., 2016; Zimmermann et al., 2017), and this needs to be researched further. In addition, the majority of studies investigating pollen chemistry have used pollen from herbaria, botanical gardens, or university campuses, with a limited number of replicates per location and large number of different species and families. Although this sampling design encourages ease of access, reliable identification, and a broad species range, the number of replicates remains a limiting factor for understanding chemical variations in response to both environmental and taxonomic factors.

This study aims to address the challenges related to understanding the taxonomic variations of pollen chemistry by investigating the relative importance of within- and between-species chemical differences in *Quercus*. Our dataset is unique because it represents the largest collection of closely related species sampled from populations outside botanic gardens across a large bioclimatic and biogeographic gradient in Portugal. We use a combined multivariate- and discriminant-analysis approach to (i) investigate the potential for FTIR as tool to differentiate six taxa of *Quercus* based on pollen, and (ii) determine the main chemical-functional groups responsible for chemical variation observed in the dataset. Addressing these questions represents the first step if we are to successfully use pollen chemistry as a tool to improve our understanding of past biogeographical patterns and history of oaks in Europe.

## 2. Methods

### 2.1 Sample collection

We collected pollen samples from 297 individual trees belonging to five *Quercus* species across a 4° (∼450 km) latitudinal gradient in Portugal (Figure 1). The *Quercus* species in this study belong to the sections *Cerris, Ilex*, and *Quercus* according to Denk et al. (2017) and have different geographic distributions (Table 1). Trees were sampled along gradients of temperature and precipitation to cover a wide range of environmental conditions. A detailed summary of the number of trees sampled at each location is in the Supplementary Material (Table S1).

**Table 1.** *Taxonomy of sampled* Quercus *trees and total number of trees sampled (n). Sections are according to Denk et al. (2017)*

| Subgenus | Section | Species | Subspecies | n | Distribution |
| --- | --- | --- | --- | --- | --- |
| Cerris | *Ilex* | *Q. coccifera* |  | 3<br>6 | *Q. coccifera* and *Q. rotundifolia* prefer xerophytic conditions and are often co-occurring species. Both are indifferent towards bedrock conditions, but prefer soils without waterlogging, although *Q. coccifera* is a thermophilous species and is less tolerant of winter cold. |
| Cerris | *Ilex* | *Q. rotundifolia* |  | 3<br>8 |  |
| Cerris | *Cerris* | *Q. suber* |  | 6<br>9 | *Q. suber* is distributed across the Mediterranean Basin on siliceous bedrock but is absent from areas with winter cold (Amigo, 2017; Matías, Abdelaziz, Godoy, & Gómez-Aparicio, 2019). |
| Quercus | *Quercus* | *Q. faginea* |  | 6<br>3 | *Q. faginea s.l.* has a broad distribution in the Iberian Peninsula (Tschan & Denk, 2012) and is more abundant on limestone with higher summer precipitation, tracking the sub-Mediterranean bioclimatic belt (Sanchez, Benito-Garzon, & Ollero, 2009). |
| Quercus | *Quercus* | *Q. robur* |  | 7<br>6 | *Q. robur* has a temperate distribution and is more abundant in north-west Portugal (Jalas & Suominen, 1976). It occurs in regions with sufficient summer rain, and is absent from areas with summer drought (Amigo, 2017; Ülker, Tavsanoglu, & Perktas, 2018). |
| Quercus | *Quercus* | *Q. robur* | *estremaduraensis* | 1<br>5 |  |

**Figure 1.**
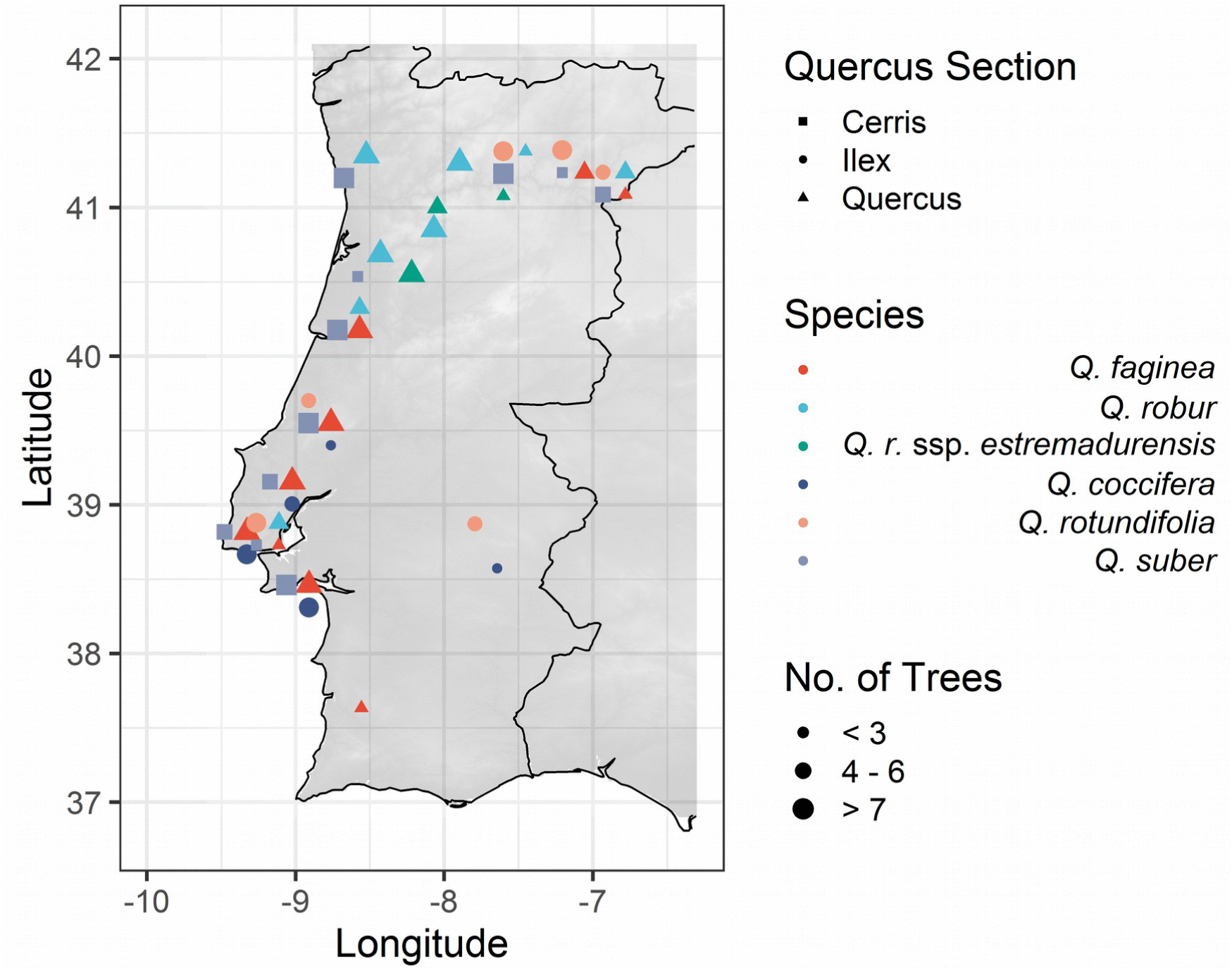
Map of Portugal displaying the locations where trees where sampled. Colour indicates species, shape indicates Quercus section following Denk et al. (2017) and size of the symbol indicates number of trees sampled per location.

All samples were collected in spring 2018 by taking whole-tree composite samples of ca. 30 catkins per individual tree. Several branches were sampled for catkins up to 5 m in height. The catkins were air-dried at room temperature (∼23 C) for at least 24h and the pollen was separated from the anthers by light shaking. Pollen was also sieved through 60 µm sieves to remove excess plant material before analysis.

### 2.2 Pollen-chemistry measurements

Reflectance-infrared spectra were recorded using a Vertex 70 FTIR spectrometer (Bruker Optik GmbH, Germany) with a single reflectance-attenuated total-reflectance (SR-ATR) accessory. The ATR IR spectra were recorded with a total of 32 scans and spectral resolution of 4 cm^−1^ over the range of 4000–600 cm^−1^, using the horizontal SR-ATR diamond prism with 45° angle of incidence on a High Temperature Golden gate ATR Mk II (Specac, United Kingdom). Approximately 1 mg of dried pollen was deposited onto the ATR crystal for each measurement (3 replicate measurements). Between each measurement a background (reference) spectrum was recorded using the sample-free setup. The OPUS software (Bruker Optik GmbH, Germany) was used for data acquisition and instrument control.

We pre-processed the spectra, since multivariate-regression methods (e.g. partial least squares; PLS) have been shown to perform better with preprocessed spectra in other studies (Zimmermann & Kohler, 2013; Woutersen et al., 2018).The processing of the spectra consisted of smoothing and calculation of the second-derivative using the Savitzky-Golay algorithm, as implemented by the EMSC package (Liland, 2017). The settings of the Savitzky-Golay smoothing algorithm (Edwards & Willson, 1974; Savitzky & Golay, 1964) were: 2^nd^ degree polynomial and a window-size of 11. The second-derivative spectra were constrained between 700 and 1900 cm^−1^ and normalized using extended multiplicative signal correction (EMSC), a MSC model extended by a linear and quadratic component (Liland, 2017). For further analyses, the mean of the measurement replicates (three) was calculated for each tree (resulting in one spectra per tree). We follow peaks of interest that have been attributed to chemical-functional groups according to Pappas et al. (2003), Gottardini et al. (2007), Schulte et al. (2008), Zimmermann (2010) and Zimmermann & Kohler (2014) (Table 2).

**Table 2.**
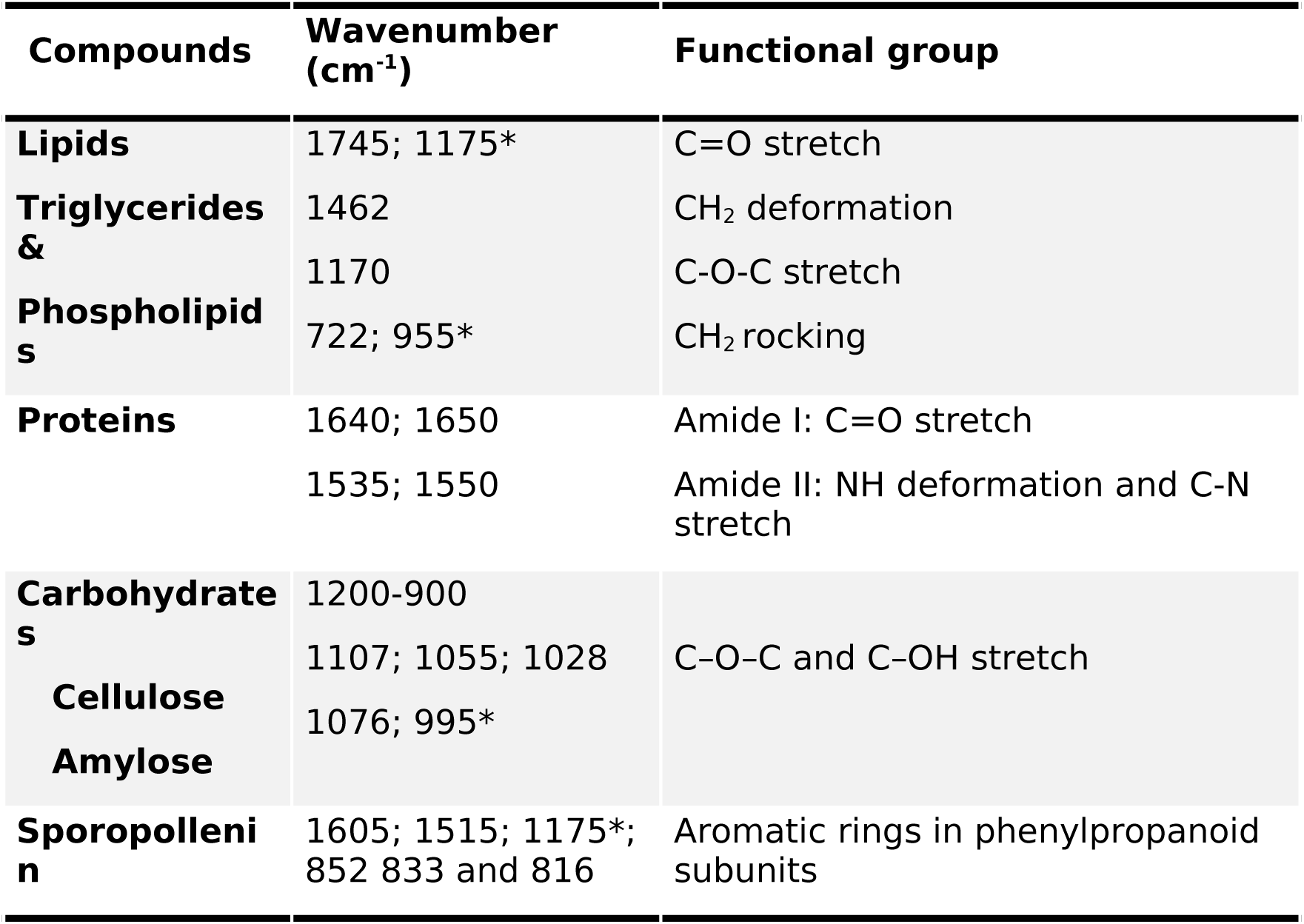
Wavenumber of peaks attributed to specific functional groups in spectra of fresh pollen and the compounds most representative for them (Pappas et al., 2003; Gottardini et al., 2007; Schulte et al., 2008; Zimmermann, 2010, p. 20109; Zimmermann & Kohler, 2014 compiled in Zimmermann, Bağcioğlu, et al., 2015). A (*) marks wavenumbers which are shared by more than one compound (Bağcioğlu et al., 2015).

### 2.3 Statistical analyses

For the exploration of within- and between-species chemical variability, we fitted a PLS model combined with canonical correlation analysis (CPPLS) to the processed mean spectra (2^nd^ derivative) to predict species identity. This analysis was implemented in the “pls” package (Mevik, Wehrens, Liland, & Hiemstra, 2019) in R version 3.6.0 (R Core Team, 2019). The PLS family of models has been shown to be powerful in multivariate analyses of FTIR-spectral data (Wold, Sjöström, & Eriksson, 2001; Liland, Mevik, Rukke, Almøy, & Isaksson, 2009; Telaar, Nürnberg, & Repsilber, 2010; Zimmermann et al., 2017). The CPPLS method improves the extraction of predictive information by estimating optimal latent variables in comparison to standard PLS regression (Mehmood & Ahmed, 2016). Unlike standard PLS, CPPLS weights the contribution of the explanatory variables (wavenumbers), which weakens the contribution of non-relevant wavenumbers to optimise the covariance between response (species) and explanatory variables (wavenumbers). Indahl, Liland, & Næs (2009) show improved accuracy and increased explained variance of CPPLS compared to conventional PLSR using spectral data.

To assess the classification performance of the CPPLS, the dataset was randomly split into training and test sets using a 60%/40% split. This split was repeated 100 times to create 100 versions of the dataset (folds) with different training/test splits. A CPPLS model was fitted for each fold and the extracted component scores were used to predict species identity using limited discriminant analyses. The performance of the classifier in predicting the test set was averaged over the folds and summarised in a confusion matrix (Table 3).

**Table 3.** *Confusion matrix of linear discriminant analysis on the test sets using 4 components of the fitted canonical powered partial least squares (CPPLS) model. Predictions as rows and Reference as columns. Values given as % of spectra that were predicted as species and sum to 100% columnwise, e.g. for Q. robur 18% of spectra were predicted as Q. faginea, 76% were correctly classified as Q. robur and 5% as estremadurensis. Abbreviations; Q. r. estr: Q. robur* ssp. *estremadurensis, Q. rotund.: Q. rotundifolia. Green signifies section Quercus, blue is section Ilex and red is section Cerris.*

| Pred/Ref | *Q. faginea* | *Q. robur* | *Q. r. estr.* | *Q. coccifer a* | *Q. rotund.* | *Q. suber* |
| --- | --- | --- | --- | --- | --- | --- |
| *Q. faginea* | 65 ± 12 | 18 ± 8 | 11 ± 13 | 0 | 0 | 1 ± 1 |
| *Q. robur* | 31 ± 12 | 76 ± 10 | 81 ± 18 | 0 | 0 | 1 ± 2 |
| *Q. r. estr.* | 1 ± 2 | 5 ± 6 | 6 ± 13 | 0 | 0 | 0 |
| *Q. coccifer a* | 2 ± 3 | 0 | 0 | 75 ± 14 | 25 ± 11 | 0 |
| *Q. rotund.* | 1 ± 2 | 0 | 0 | 25 ± 14 | 75 ± 11 | 0 |
| *Q. suber* | 3 ± 4 | 1 ± 3 | 2 ± 6 | 0 | 0 | 99 ± 2 |

## 3. Results

### 3.1 Chemical variations in *Quercus*

Assessment of the mean spectra of the five *Quercus* species and one subspecies reveals clear differences in absorbance between the major intrageneric lineages (sections) at wavelengths associated with specific chemical functional groups (Figure 2). For example, the lipid peak absorbance at ∼1750 cm^−1^ is weaker in section *Ilex* compared to the other sections, whilst the sporopollenin and carbohydrate absorbance bands (at 1516, 1168, 833 cm^−1^ and 985 cm^−1^, respectively) are noticeably lower in absorbance in the taxa belonging to section *Ilex*. Note, however, that although clear spectral differences exist at the section level, it is more difficult to separate variations between species *within* different sections (Figure 2).

**Figure 2.**
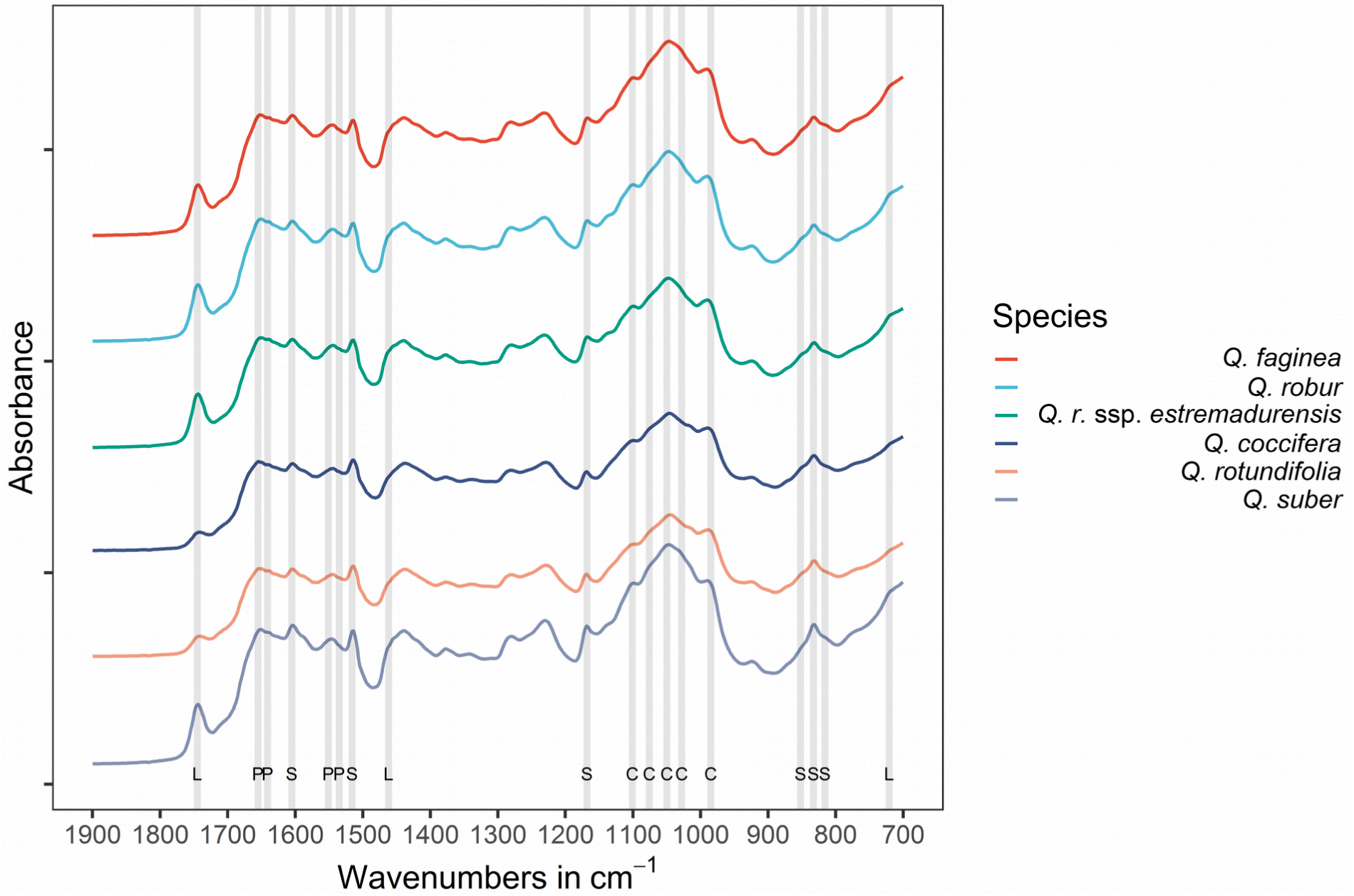
*Mean absorbance spectra of* Quercus *species and notable peak locations. Lipids (L), protein (P), sporopollenin (S), and carbohydrates (C) using wavenumbers given in Table 2. Spectra are offset.*

The observations made following assessment of the mean spectra are confirmed by the analysis using CPPLS (Figure 3a). Here, the three sections of *Quercus* can be clearly separated using the variance explained by two CPPLS components (Figure 3a). For example, the section *Ilex* scores are negative along the first component (26 % of the variation), whilst individuals of sections *Quercus* and *Cerris* have positive scores along this axis. The sections *Quercus* and *Cerris* can mainly be separated along the second principal component (section *Cerris* with positive scores and section *Quercus* with negative scores). However, between-species level variations within species in the same section are harder to differentiate, because there is a large overlap between species of the sections *Ilex* and *Quercus*. This overlap is reduced on later components, where the different species separate within their respective sections. On components 3 and 4 *Q. coccifera* and *Q. rotundifolia* can be separated along the third component, while *Q. robur* and *Q. faginea* show some separation along the fourth component (Figure 3b). In total, the two first components explain ∼37% of the variation in the dataset and separate the samples into the *Quercus* sections, while the next two components explain a further ∼9.7% (Figure S1).

**Figure 3.**
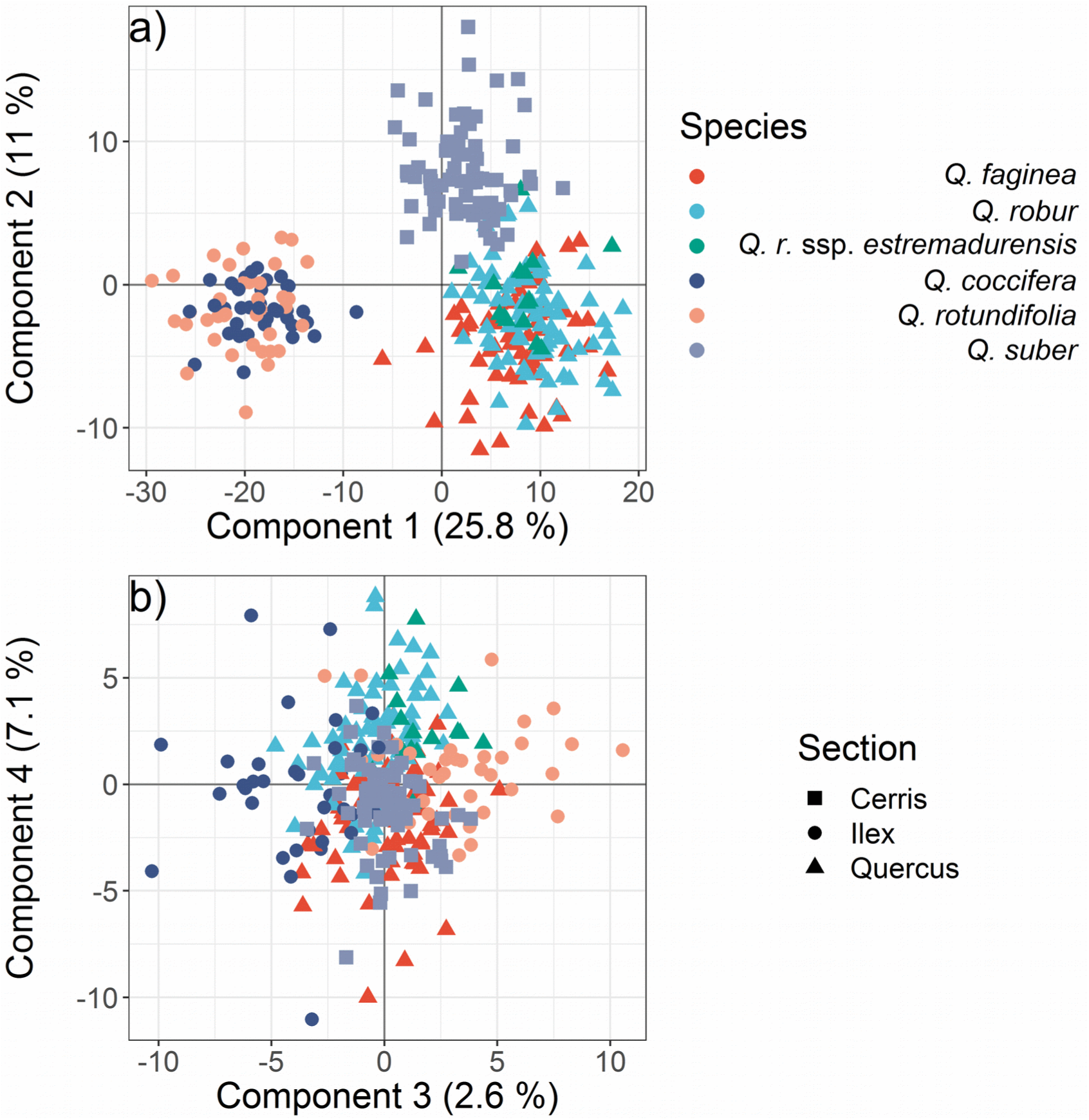
*Ordination of components 1-4 of canonical powered partial least squares (CPPLS) models (100-fold cross-validation). Each point represents a sampled tree. The mean training scores for each sample over the 100-folds were calculated. Proportion of variance explained by each component in parentheses. Colour indicates species, while symbol indicates* Quercus *section*.

Since specific absorbance peaks in the FTIR spectra can be related to chemical functional groups (summarised in Table 2), it is possible to identify which chemical functional groups have the highest influence in explaining patterns of variation observed in our dataset (Figure 4). In general, lipids have the highest loadings (i.e. greatest influence) on component 1. The peak at 995 cm^−1^ in pollen can also be attributed to phospholipids (Bağcıoğlu et al., 2015) in addition to carbohydrates. Sporopollenin peaks have high positive loadings on component 1 and mostly negative loadings on component 2. The 852 cm^−1^ sporopollenin peak shows high positive loadings on component 2 in contrast to the other sporopollenin peaks (816, 833, 1175, 1516, and 1605 cm^−1^). Components 3 and 4 have higher loadings for carbohydrates and protein peaks. Taken together, these results indicate that sporopollenin and lipids are strong drivers of the main sources of variation in the dataset and congeneric discrimination is achievable based on FTIR at least at section level.

**Figure 4.**
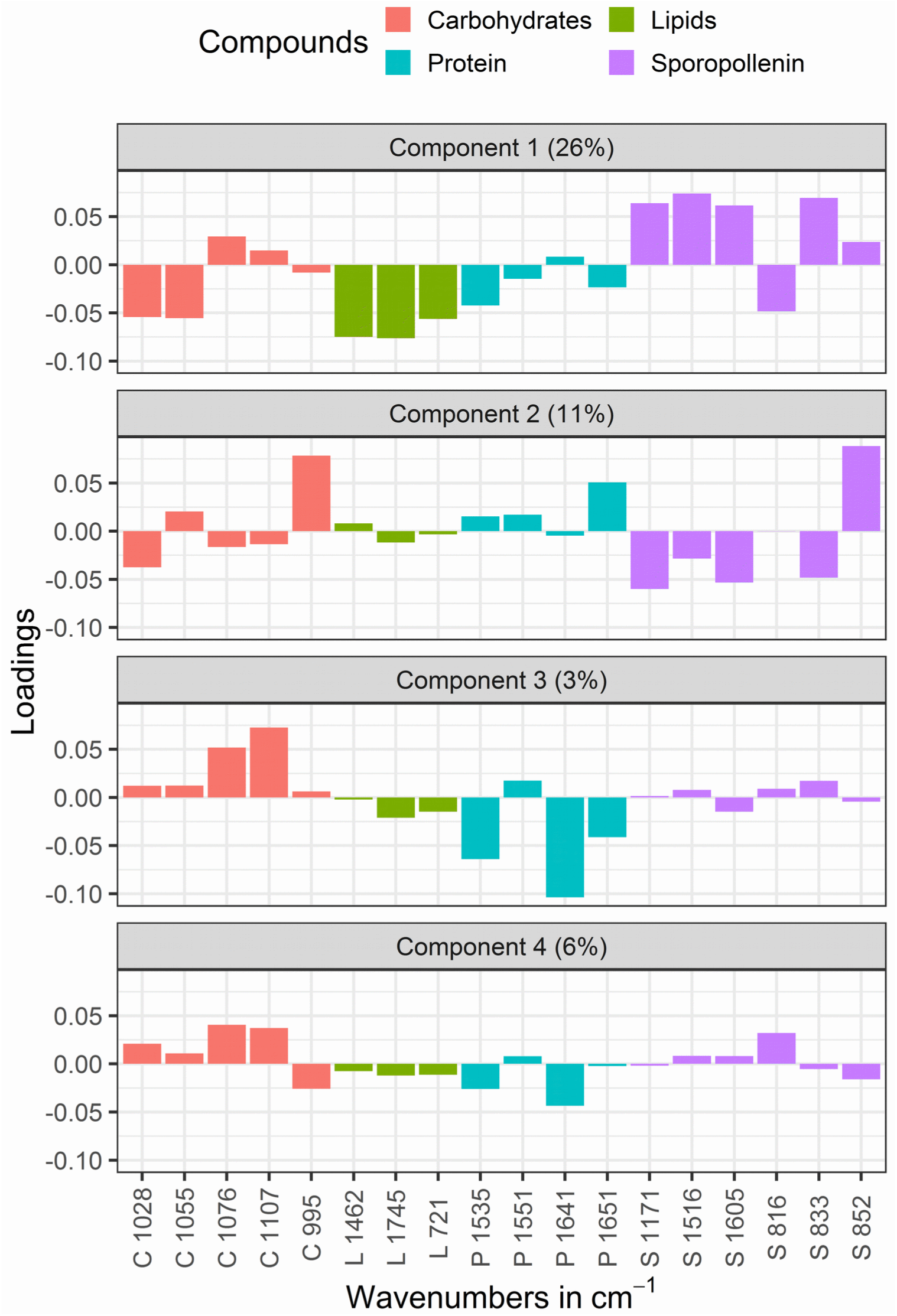
Loadings plot of classification of canonical powered partial least squares (CPPLS) model. Lipids (L), protein (P), sporopollenin (S), and carbohydrates (C) using wavenumbers given in Table 2. High absolute loading indicates a high importance of a given wavenumber for the corresponding component. Loadings are chosen in such a way as to describe as much as possible of the covariance between the variables (wavenumbers) and the response (species). Proportion of variance explained by each component in parentheses.

### 3.2 Discriminant analysis

Using four components (explaining ∼45% of the variance) the confusion matrix of the classification CPLS model shows clear differentiation between the *Quercus* sections, with some misidentified spectra (<∼2 ±3) from section *Quercus* and *Q. suber* (Table 3). *Quercus robur* ssp. *estremadurensis* has by far the worst accuracy in the model and is most often identified as its parent species *Quercus robur*, possibly due to the limited number of samples in the dataset (Table 1). In general, species misidentifications are contained within the different *Quercus* sections and lie between 18% and 31% of the test-set samples (within-section misidentifications: 18% of *Q. robur* as *Q. faginea*; 31% of *Q. faginea* as *Q. robur*; 25% of *Q. coccifera* as *Q. rotundifolia*; 25% of *Q. rotundifolia* as *Q. coccifera*). Overall species accuracy within sections ranges from 65% to 76% in the Ilex and Quercus sections. Increasing the components available to the model to 10 (∼57% explained variance) increases species accuracy by 5-10 percentage points in both sections (*Quercus, Ilex*) (Table S2). As demonstrated with our ordination plots (Figure 3), differentiation of the three sections of *Quercus* is possible using 37% of the variance in the spectral data, but these results indicate difficulties in differentiation between species of the same section.

## 4. DISCUSSION

### 4.1 Separation of *Quercus* according to chemical variation

Recent research has shown that spectroscopic methods such as FTIR are effective at differentiating pollen species between distantly related families and/or genera using their chemical composition (Gottardini et al., 2007; Dell’Anna et al., 2009; Zimmermann, 2010; Zimmermann & Kohler, 2014; Zimmermann, Bağcıoğlu, et al., 2015; Zimmermann, Tafintseva, Bağcioğlu, Berdahl, & Kohler, 2016; Julier et al., 2016; Zimmermann et al., 2017; Woutersen et al., 2018; Jardine et al., 2019). Our results build on these previous studies to reveal the potential for chemical variations in pollen to distinguish infrageneric variation between species in pollen samples from 297 individuals from Portugal belonging to three different *Quercus* sections (*Cerris, Ilex, Quercus*). We identify a clear separation at the *Quercus* section level (Fig. 3 and Table 3), using two components of a PLS model and 37% of the explained variance in the spectral data. One component (component 1) can be used to differentiate the sections *Ilex* and *Quercus*, while component 2 can be used to separate section *Cerris*. Combined, these two components achieve the performance equivalent to SEM methods, where *Quercus* pollen can be confidently identified to section level.

Despite finding that classification at the section level is possible using FTIR approaches, there is considerable overlap in variation between species of the same section. Furthermore, classification performance does not improve when using a more complex model in which the number of components used increases from four to ten. Using this more complex model, which explains ∼57% of the variance (compared to ∼45% in the four-component model), classification accuracies remain roughly similar within *Quercus* sections (Table S2). For example, *Q. coccifera* and *Q. rotundifolia* have a recall of ∼75% with both 4 and 10 components. Similarly, approximately one third of *Q. robur* and *Q. faginea* samples (both belonging to section *Quercus*) are misclassified as the other species. Thus, while our results indicate that subgeneric classification of *Quercus* pollen is possible at the section level using FTIR, we still find it difficult to distinguish between more closely related (i.e. within-section) pollen types.

These findings are approximately in line with other studies, which have performed species classification using FTIR. For example, both Julier et al. (2016) and Jardine et al. (2019) report classification successes of ∼80% and ∼85% rates, respectively, using an FTIR analysis of cryptic morphospecies within the family Poaceae. Their studies are based on a combination of specimens of mainly non-congeneric grass species (except two species of *Oryza*, Julier et al., 2016, and four species of *Triticum*, Jardine et al. 2019). In both these studies, classification success is lower for the samples belonging to congeneric species and higher for the more-distantly related pollen types (i.e. those species belonging to different genera). In another study, Woutersen et al. (2018) report ∼95% recall on largely congeneric species in the Nitrariaceae family using single grain FTIR, but also note that lack of environmental variability (pollen from one individual per species) could have led to an overestimation of classification success. In contrast, Zimmermann et al. (2017) achieve ∼100% accuracy on species identification and 75% accuracy on identification of origin using hierarchical PLSR on pollen from three species of Poaceae (*Festuca ovina, Anthoxanthum odoratum, Poa alpina*) of different genera and origins (Sweden, Norway, Finland) grown under controlled conditions (45 individuals per species). Such a high classification success on taxa grown in controlled conditions demonstrates the strong phylogenetic signature that can be observed using FTIR. Our results also demonstrate these strong phylogenetic differences in FTIR spectra (i.e. the ability to differentiate between *Quercus*-section level variability), but our study additionally demonstrates the difficulty of distinguishing between-species level variability, even when relatively large subsets of samples are used.

### 4.2 Key chemical drivers of variation within *Quercus* spp. pollen

Given the result that identification is possible at the section level, a key question that follows is which of the chemical components of the pollen grain are mostly responsible for the difference between the three sections of *Quercus* under FTIR? In our study, we find that lipids are one of the most important functional groups in diagnosing samples belonging to section *Ilex*. The waveband at 1750 cm^−1^ is particularly important in this regard. Previous research has shown that this waveband is a strong indicator for triglyceride lipids (Bağcioğlu et al., 2017). Our results also confirm previous findings by Zimmermann & Kohler (2014), who show extreme variations in the relative content of triglycerides and find this waveband to be an important separator between *Iris, Quercus*, and *Pinus* pollen types. Indeed, our results extend the inferences made in that previous study by demonstrating a subgeneric level variability of the relative lipid content. Specifically, we identify a distinctly lower amount of lipids in pollen sampled from individuals within the Ilex section (Figure 2).

In addition to the importance of triglyceride lipids as a tool for chemical separation, we also find that wavebands representing building blocks of sporopollenin (Table 3) are important for differentiating taxa on the first two components of our CPPLS analysis. For example, the peaks at 833, 852, 1516 and 1605 cm^−1^ are associated with building blocks of sporopollenin (Bağcioğlu et al., 2015) and have the highest loadings on component 2, which is used in this study to isolate *Q. suber*. Peaks at 833 and 852 cm^−1^ are related to different types of phenylpropanoid building blocks and our results suggest relative differences in their abundance within the sporopollenin of *Q. suber* compared to the other species. In addition to lipid variation, aromatic peaks at 833, 852, 1516 and 1605 cm^−1^ also have a high loading on component 1, which can be used to separate the *Ilex* section pollen from the other sections. Thus, both lipids and sporopollenins are important functional groups to different pollen between the three sections in our dataset.

Our observations of different chemical compositions of sporopollenin mirror the sequence of the development of the pollen wall in different *Quercus* sections described by Solomon, (1983a, 1983b) and Denk & Grimm, (2009). Evolutionary, pollen of section *Ilex* represent the earliest, primitive state of *Quercus* pollen, with a microrugulate pattern on the pollen exine surface. A set of secondary sporopollenins are then added to this surface during exine formation in pollen of sections *Cerris* and *Quercus* (Denk & Grimm, 2009). It is possible that these key differences in structure and formation of the exine between the section *Ilex* and other *Quercus* pollen grains are responsible for the key differences sporopollenin chemistry identified using FTIR. More detailed work on the composition of sporopollenin of different genera and how this affects pollen grain structural elements (e.g. Li, Phyo, Jacobowitz, Hong, & Weng, 2019) is needed for this finding to be confirmed.

Finally, protein and carbohydrate peaks (Carbohydrates: 1107, 1028, 1076 cm^−1^; Proteins: 1535, 1641 cm^−1^) have the highest loadings on components 3 and 4 and are partly responsible for the partial distinction of species within the same section. These peaks represent amylose and cellulose as carbohydrates and amide functional groups within proteins (Table 2). For example, variation along component 3 contributes to the separation of *Q. robur* from *Q. faginea*. However, overall, protein and carbohydrate peaks have the least influence for explaining the variance of classification success, and most of the taxonomically important information we used to distinguish between the species is stored in the lipids and sporopollenin components of the pollen chemistry.

### 4.3 Implications for understanding past and future *Quercus* dynamics

We investigated the potential for chemical separation of *Quercus* pollen because, despite the high diversity (22 species) of *Quercus* in Europe, fossil pollen of this genus is still most commonly determined as either *Quercus* Deciduous or *Quercus* Evergreen types (e.g. Huntley & Birks, 1983; Brewer et al., 2017). As a result, there may be much detail missing in our current understanding about past *Quercus* dynamics which could be improved through methods that result in refined taxonomic resolution. Indeed, our results indicate the potential for FTIR to surpass traditional LM methods used in palynology, and work at a comparable level to SEM (Denk & Grimm, 2009; Denk & Tekleva, 2014; Grímsson et al., 2015; Grímsson, Grimm, Meller, Bouchal, & Zetter, 2016). However, the extensive automated-classification possibilities offered by future IR analysis (Mondol et al., 2019), and in the ease of sample preparation and data collection, may mean it will be easier to expand these technologies compared to the more time consuming SEM methods in the long-term.

The ability to differentiate at higher taxonomic resolution would enhance our understanding of past trajectories of co-occurring *Quercus* sections, in particular for understanding the expansion of *Quercus* since the Last Glacial Maximum (e.g. S. Brewer, Cheddadi, de Beaulieu, & Reille, 2002). FTIR techniques may also be useful for older interglacial sequences, where identification of *Quercus* pollen to section is often not possible due to degradation (Tzedakis, 1994). This would complement studies that use genetic methods to reconstruct colonization pathways, which have higher taxonomic resolution and compliment the palynological data, but lack the temporal resolution that pollen records provide (Petit et al., 2002).

In addition, a number of studies have highlighted the need to incorporate long-term ecological information to improve biodiversity forecasts of environmental change (Dawson, Jackson, House, Prentice, & Mace, 2011). Rates of temperature increases in the Mediterranean are projected to outpace the rest of the temperate regions in Europe and are predicted to rapidly change the associated biomes in the region (Giorgi & Lionello, 2008; Guiot & Kaniewski, 2015; Guiot & Cramer, 2016), but the consequences of this change for Mediterranean oak forests remain uncertain (Lindner et al., 2014; Acácio et al., 2017). A number of studies have integrated pollen data in order to reduce uncertainties when forecasting the biotic responses of *Quercus* to climate change in the future (Nogues Bravo et al. 2016; Guiot & Cramer, 2016). However, like the palaeoecological studies discussed above, the limited taxonomic resolution used may bias projections. For example, species distribution models based solely on *Quercus* pollen, were only able to estimate niche-environment relationships at the genus level. Extensive application of FTIR techniques may therefore provide a bridge between long-term ecological and modern biogeographic approaches.

Nevertheless, despite the potential shown in our FTIR approach, our findings still reveal a number of challenges before vibrational methods can be rolled out across biogeographic and palaeoecological applications. First, our results are still unable to resolve at the species level, and so although taxonomic resolution would be refined using FTIR approaches, in many cases palaeoecological studies would still lack the taxonomic precision of other biogeographic tools (e.g. phylogenetic analysis). Second, our results are based on fresh pollen sampled from modern taxa, and lipids were the main functional groups used to differentiate between taxa in this study, followed by sporopollenins (Figure 4). Although the preservation and stability of sporopollenins in fossil sequences are well established (Fraser et al., 2012), the extent to which lipids are preserved in chemical sequences in subfossil pollen sequences remains uncertain. Variations in sporopollenin functional groups were still responsible for differentiation between the three main *Quercus* sections, but the ability for these functional compounds to be used as taxonomic tools in isolation is yet to be established. In the future it may be more beneficial to focus on variations of sporopollenins in pollen, perhaps through the use of Raman spectroscopy, which preferentially targets the vibration of non-polar bonds in sporopollenins and so may be able to achieve finer-scale differentiation of sporopollenin building blocks (Merlin, 2009).

Third, in this study we used bulk pollen samples to infer differences using FTIR, but fossil-pollen samples would require single-grain measurements since pollen grains are difficult to separate from other organic material within the sediment matrix. Single-grain FTIR spectra are less reproducible than bulk, mostly due to spectral anomalies caused by scattering and by non-radial symmetry of certain pollen types (Zimmermann, Bağcioğlu, et al., 2015; Zimmermann, 2018). Although these issues have been addressed by adjusting experimental settings and by implementing numerical correction methods (Zimmermann et al., 2016; Zimmermann, 2018), future work is needed to test whether the patterns we observe at the bulk level can be replicated using single-grain FTIR measurements.

Finally, our study shows the importance of using large numbers of replicates in the pollen samples to account for the large amounts of chemical variation present in the chemical spectra, even within replicate species. The large numbers of samples and high levels of replication here (i.e. 50 ± 23 tree replicates per species) are a major advantage over previous studies, which have featured either fewer replicates (<5)(Jardine et al., 2019; Julier et al., 2016; Woutersen et al., 2018) or fewer/no congeneric species (Julier et al., 2016; Zimmermann et al., 2017). Although we do find clear signals in the data linked to systematics (Figure 3 and Table 3), we also find that ∼60 % of the total variation remains unexplained. One probable reason for the unexplained variation observed in our study may be linked to the environmental controls on pollen chemistry. Previous studies have suggested plasticity of pollen chemistry to climate and other environmental variables (Zimmermann & Kohler, 2014; Depciuch et al., 2016; Bağcıoğlu et al., 2017; Depciuch, Kasprzyk, Sadik, & Parlińska-Wojtan, 2017; Zimmermann et al., 2017). The other probable reason is the intra-species variation between the genotypes of different populations as well as within populations (Zimmermann et al., 2017). This suggests it will be critical to understand the other factors which can account for this variation if these pollen-chemistry techniques can be successfully applied to fossil sequences.

## 5. Conclusions

We investigated the chemical variation in pollen sampled from 297 individuals of *Quercus* using FTIR to investigate whether this technique could enable taxonomic discrimination of modern *Quercus* pollen. Our results achieved excellent (∼97%) recall to section level, showing that subgenus level differentiation of pollen samples is possible using IR methods. However, despite these promising results at the section level, more detailed, species-level differentiation was complicated by overlapping variation in the chemical composition of closely related species.

We also aimed to identify whether variations in specific functional groups are responsible for any taxonomic discrimination in the data. Here, we found lipids and sporopollenins to be key determinants between different *Quercus* sections. Although the sporopollenin functional groups are identified as important for discrimination between *Quercus* taxa, isolating the effect of these sporopollenin groups from the effects of other functional groups which may not be preserved in sediment sequences (e.g. lipids) still present a challenge. In addition, testing the application on single-grain *Quercus* samples, and developing a more complete understanding of the effects of environmental variation on pollen-chemical signatures in *Quercus* is required. Taken together, our findings build on previous studies and show that, whilst FTIR approaches on modern *Quercus* pollen can perform at a similar level to SEM techniques, future work on the discrimination of sporopollenin components is required before FTIR can become a more widespread tool in long-term ecology and biogeography. Thus, our study represents a valuable step forward in improving our understanding of variation in pollen chemical composition and its application in long-term ecology and biogeography.

## v. Acknowledgements

We dedicate this paper to the memory of John Flenley (1936-2018) who pioneered so many exciting and novel aspects of pollen analysis and vegetation history. One of us (HJBB) knew John since 1963 and was always stimulated by John’s latest ideas and palaeoecological studies. John would have been fascinated by the potential of pollen chemistry in palaeoecology and biogeography.

## x. Data Availability

Data and code for analysis are available as supplementary material on a github repository (https://github.com/FM-uib/Quercus_Portugal_FTIR) for review. Both will be uploaded to a dryad repository.

## xii. Biosketch

Florian Muthreich is a palaeoecologist interested in developing new methods of pollen classification. This work represents a component of his PhD work at the University of Bergen University within the PollChem project (https://www.uib.no/en/rg/EECRG/98775/pollchem). In this project he and other authors collaborate to explore pollen chemistry applications in biogeography and long-term ecology.

## Author contributions

FM, AWRS, BZ and HJBB conceived the idea; FM and CMVV conducted the fieldwork and collected the data; FM, BZ, and AWRS analysed the data; and FM and AWRS led the writing with assistance from BZ, HJBB, and CMVV

