## Supplementary Material for "Chemical variations in *Quercus* pollen as a tool for taxonomic identification: implications for long-term ecological and biogeographical research"

### xiv. Supporting Information

#### Taxonomic scheme of studied *Quercus* species

**_Quercus_ Linnaeus (1753 : 994)**

**I. _Quercus_ subgen. Cerris Oersted (1866: 74)**

**i. Section Cerris Dumortier (1829: 15)**

**1. _Quercus_ suber L.**

**iii. Section Ilex Loudon (1838: 1899)**

**2. _Quercus coccifera_ L. (1753: 995)**

**3. _Quercus rotundifolia_ Lam.**

**II. _Quercus_ subgen. Quercus Hickel & Camus (1921: 379)**

**vi. Section Quercus Linnaeus (1753: 994) (≡Section Robur Loudon (1838: 1731))**

**4. _Quercus_ robur L. (1785: 725)**

**iv. subsection Galliferae (Spach) Guerke**

**5. _Quercus_ faginea Lam.**

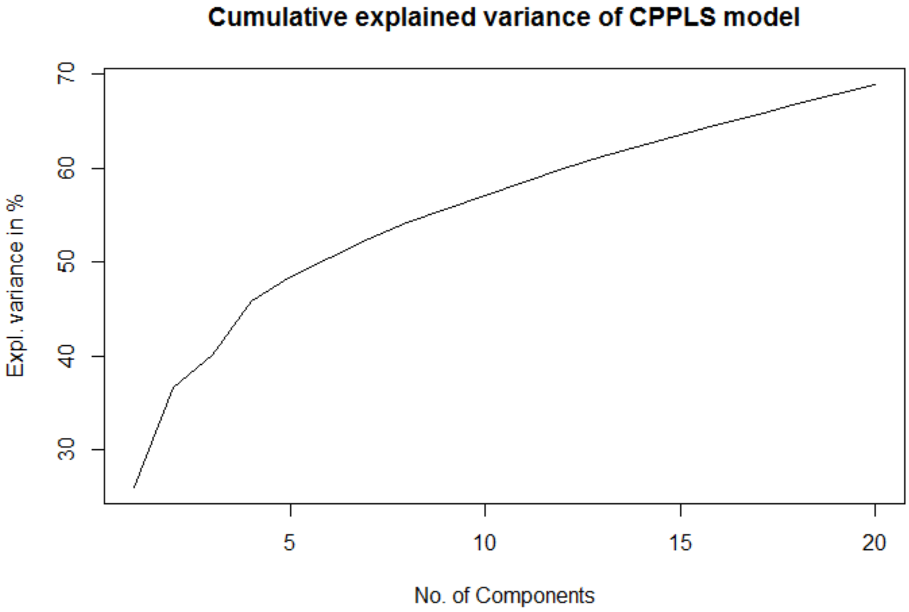


*Figure S1 Cumulative explained variance for Classification CPPLS*

*Table S1 Summary statistics of precipitation and temperature in the sampled locations in month of sampling. (April 2018). Number of sampled trees at each location.*

| | Precipitation | | | Temperature | | | | | Species | | | | | |
|---|---|---|---|---|---|---|---|---|---|---|---|---|---|---|
| Group | mean | min | max | mean | min | max | Lat. | Lon. | Q. fag. | Q. rob. | Q. r. est. | Q. coc. | Q. rot. | Q. sub. |
| Peso da Regua | 17 | 3 | 38 | 11.1 | 9.0 | 14.9 | 41.151 | -7.972 | | 15 | 6 | | | |
| Arrabida | 3 | 1 | 21 | 16.2 | 13.9 | 19.9 | 38.486 | -9.011 | 18 | | | 20 | | 9 |
| Alijo | 25 | 18 | 34 | 12.5 | 11.1 | 15.7 | 41.228 | -7.527 | | 2 | 1 | | 7 | 9 |
| Horta da Vilarica | 20 | 10 | 26 | 13.2 | 11.3 | 15.1 | 41.235 | -7.131 | 5 | | | | 9 | 3 |
| Candeeiros | 68 | 30 | 116 | 14.1 | 13.1 | 15.0 | 39.550 | -8.838 | 7 | | | 3 | 5 | 8 |
| Freixo de Espada A Cinta | 16 | 9 | 31 | 10.5 | 8.4 | 13.5 | 41.087 | -6.856 | 3 | 4 | | | 5 | 4 |
| Porto | 15 | 5 | 25 | 13.8 | 10.8 | 16.5 | 41.199 | -8.599 | | 27 | | | | 13 |
| Coimbra | 25 | 8 | 46 | 14.9 | 13.6 | 15.6 | 40.175 | -8.644 | 7 | 5 | | | | 8 |
| Sintra | 14 | 11 | 25 | 16.9 | 14.5 | 17.6 | 38.743 | -9.428 | 8 | | | 8 | | 5 |
| Odemira | 10 | 10 | 10 | 15.0 | 14.9 | 15.1 | 37.633 | -8.633 | 2 | | | | | |
| Oliveira do Bairro | 103 | 32 | 128 | 13.7 | 12.9 | 16.3 | 40.536 | -8.506 | | 9 | | | | 3 |
| Sao Pedro do Sul | 78 | 40 | 166 | 11.2 | 10.6 | 12.0 | 40.702 | -8.145 | | 9 | 8 | | | |
| Montejunto | 90 | 18 | 120 | 14.8 | 14.2 | 16.0 | 39.180 | -9.037 | 8 | | | 4 | | 4 |
| Evora | 48 | 31 | 62 | 10.5 | 9.3 | 11.0 | 38.723 | -7.718 | | | | 1 | 5 | |
| Lisbon | 1 | 0 | 5 | 16.7 | 15.7 | 19.7 | 38.729 | -9.188 | 2 | 5 | | | 7 | 3 |

*Table S2 Confusion Matrix of linear discriminant analysis on the test sets using 4 components of the fitted canonical powered partial least squares CPPLS model. Predictions as rows and Reference as colums. Values given as % of counts that were predicted as species. Sum to 100 % columnwise, e.g. For Q. robur 17% of spectra were predicted as Q. faginea, 76% were correctly classified as Q. robur and 7% as estremadurensis. Q. r. estr: Q. robur estremadurensis, Q. rotund. :Q. rotundifolia. Green signifies Quercus section, blue is Ilex section and red is Cerris section*

| ***Pred/Ref*** | ***Q. faginea*** | ***Q. robur*** | ***Q. r. estr.*** | ***Q. coccifera*** | ***Q. rotund.*** | ***Q. suber*** |
|---|---|---|---|---|---|---|
| ***Q. faginea*** | 74 ±10 | 17 ±7 | 6 ±9 | 0 | 0 | 0 |
| ***Q. robur*** | 19 ±8 | 74 ±9 | 77 ±22 | 0 | 0 | 0 |
| ***Q. r. estr.*** | 4 ±5 | 9 ±7 | 16 ±19 | 0 | 0 | 0 |
| ***Q. coccifera*** | 0 | 0 | 0 | 77 ±11 | 26 ±12 | 0 |
| ***Q. rotund.*** | 0 | 0 | 0 | 23 ±12 | 74 ±12 | 0 |
| ***Q. suber*** | 2 ±2 | 0 | 0 | 0 | 0 | 100 ±1 |
